# Inhibition of Quorum Sensing System in *Pseudomonas aeruginosa* by Psammaplin A and Bisaprasin Isolated from the Marine Sponge *Aplysinella rhax*

**DOI:** 10.1101/2021.01.22.427779

**Authors:** Emmanuel T. Oluwabusola, Nursheena Parveen Katermeran, Lik Tong Tan, Oluwatofunmilayo Diyaolu, Jioji Tabudravu, Rainer Ebel, Marcel Jaspars

## Abstract

Natural products isolated from marine sponges have exhibited profound bioactivity and, in some cases, serve as potent quorum sensing inhibitory agents by preventing microbial biofilm formation. In this study, the inhibitory potential of the psammaplin type compounds, psammaplin A (**1**) and bisaprasin (**2**), isolated from the marine sponge, *Aplysinella rhax*, was evaluated in the quorum-sensing inhibitory assay based on the *Pseudomonas aeruginosa* PAO1 *lasB-gfp* and *P. aeruginosa* PAO1 *rhlA-gfp* biosensor strains. The result indicated that psammaplin A (**1**) showed moderate inhibition against *lasB-gfp* biosensor strains but significantly inhibited the QS-gene promoters, *rhlA-gfp* with IC_50_ value at 30.69 and 2.64 μM, respectively. In contrast, bisaprasin (**2**) displayed significant inhibition for both biosensor strains, *lasB-gfp* and *rhlA-gfp* with IC_50_ values at 8.70 and 8.53 μM, respectively. To our knowledge, the antiquorum sensing activity of these marine-derived bromotyrosine compounds is described here for the first time.

**Significance and Impact of the Study:** The attention of the scientific community has been drawn to using marine sources to find novel quorum sensing inhibitors as antipathogenic drugs to combat antimicrobial resistance and pathogenesis caused by the proliferation of pathogenic bacteria mediated by the quorum sensing (QS) system. By blocking the QS signalling communication, the ability to assemble an organised community structure that enables drug resistance and production of virulence factors will be attenuated. The significance of this investigation is based on the discovery of bromotyrosine derivatives as potential new drug leads for the development of antipathogenic agents.

## Introduction

The discovery of antibiotics in the early 20th century was life-saving for people suffering from infectious diseases(Aminov *et al*. 2010). Despite the progress made in drug development, there is a high demand for better strategies to tackle the widespread threat of antibiotic-resistant microorganisms. Unfortunately, many common community-acquired pathogenic bacteria, such as *Escherichia coli, Klebsiella pneumoniae, Salmonella*, methicillin-resistant *Staphylococcus aureus* and *Pseudomonas aeruginosa*, have shown progressive resistance to conventional drugs. A World Health Organisation (WHO) report in 2019 concluded that if the current trend is not averted drug-resistant diseases could lead to the death of 10 million people each year by 2050 (Raymond and Powell 2019; Dai *et al*. 2019).

Intensive research in bacterial cellular functions implicates the quorum sensing system as the major contributor to their continual pathogenesis and drug resistance. Quorum sensing (QS) is a cell-to-cell communication system exhibited by most microorganisms in response to their cell density (Ng and Bassler, 2009). The QS system involves the secretion of chemical signalling molecules to monitor population density within bacterial communities. As the bacterial colony size increases, it generates a sufficient number of signal molecules to activate a variety of downstream cellular processes, including biofilm formation, virulence factors, and drug resistance mechanisms (Zhao *et al*. 2020). QS signalling of pathogens represents the central regulatory mechanism for the expression of virulence determinants, such as proteases, elastases, exotoxins, rhamnolipids and many other immune-evasion molecules (Pamp and Tolker-Nielsen 2007; Chen *et al*. 2013). For instance, *P. aeruginosa* possesses four functional QS systems consisting of LasI/LasR, Rhll/RhlR pseudomonas quinolone signal (PQS) and the integrated quorum-sensing signal (IQS) (Davies 1998; Jakobsen *et al*. 2013). LasI/LasR and Rhll/RhlR circuit homologs are the two acyl homoserine lactone (AHL)-dependent QS system in *P. aeruginosa* (Wagner *et al*. 2006). The interaction of AHLs, namely *N*-3-oxododecanoyl homoserine lactone and *N*-butyryl-homoserine lactone, with the respective receptors, lasR and rhIR, activates the transcription of nearly 10% of about 300 genes in *P. aeruginosa*. (Schuster and Peter, 2006). In addition, cell-to-cell signaling controlled the expression of *lasB and rhlA* genes, which encodes for the major virulence factor, elastase and rhamnolipids, respectively (Pesci *et al*. 1999; Rasmussen and Givskov, 2006).

Studies have shown that biofilm forming microorganisms exhibit far more resistance to antibiotics than their planktonic counterparts (Rutherford and Bassler 2012). Rhamnolipids are amphipathic glycolipids and are produced through the *rhlAB* operon and *rhlC*. These compounds plays multiple functions in the maturation and preservation of biofilms by assisting in the formation of microcolony and extracellular polymeric substances (EPS) that are embedded in the bacterial community (Rahim *et al*. 2001; Davey *et al*. 2003; Pamp and Tolker-Nielsen, 2007). The biofilm mode of development serves as a survival strategy for pathogenic microorganism to increase antibacterial resistance and cause severe systemic infections (Chen *et al*. 2018). A study conducted by Davies and co-workers revealed a correlation between signalling molecules and biofilm development in *Pseudomonas aeruginosa* and has paved the way for further biofilm research (Davies 1998). As such, interference of the QS system in pathogenic bacteria represents an attractive target for the development of novel therapeutics. For instance, the use of small molecules as potential inhibitors against QS mediated genes responsible for virulence determinants and biofilm formation is an effective strategy in combating microbial resistance to conventional antibiotics (Dai *et al*. 2019).

Several marine-derived natural products have shown considerable potential as quorum sensing inhibitors, for example manoalide, secomanoalide and manoalide monoacetateisolated from the marine sponge, *Luffariella variabilis*, exhibited potent QS inhibition against *lasB*-*gfp* (ASV) biosensor with IC_50_ values at 0.66 μM, 1.11 μM and 1.12 μM, respectively (Skindersoe *el al*. 2007). In addition, aplyzanzines C–F, a series of bromotyrosine alkaloids isolated from the French Polynesian sponge *Pseudoceratina n*. sp., showed quorum sensing inhibition against the wild typed *V. harveyi* strain (Tintillier *et al*. 2020).

In our search from marine sources for novel quorum sensing antagonists against QS *lasB* and *rhlA* genes promoters responsible for virulence factors and biofilm formation in *P. aeuroginosa*, we tested psammaplin **A** (**1**) and bisaprasin (**2**) (Rodriguez *et al*. 1987; Jimenez and Crews, 1991; Park *et al*., 2003). The compounds were previously isolated alongside psammaplins **B**-**D** (Rodriguez *et al*. 1987; Jimenez and Crews, 1991; Park *et al*., 2003) and **O**-**P**, 3-bromo-2-hydroxyl-6-carbomethoxy benzoic acid (Oluwabusola *et al*. 2020), 2-(3-bromo-4-hydroxyphenyl) aceto-nitrile and 3-bromo-4-hydroxybenzoic acid from the methanolic extract of the marine sponge, *Aplysinella rhax*, collected from the Fiji islands and were subjected to QS inhibitory screening. Psammaplin A (**1**) is composed of two modified amino acids: a bromotyrosine containing an oxime group, and cysteamines that form the disulfide bridge (Rodriguez *et al*. 1987). The bromotyrosine compound (**1)** and its derivatives have shown to possess good cytotoxicity against ovarian tumour cell lines (Tabudravu *et al*. 2002), considerable antibacterial activity against gram-positive methicillin-resistant *S. aureus*, inhibition of DNA synthesis and the supercoiling activity of DNA gyrase (Kim *et al*. 1999), and antiparasitic activity (Oluwabusola *et al*. 2020). The current study was conducted against the backdrop of interesting biological activities of these type of bromotyrosine compounds. Herein we report for the first time the inhibitory activity against the QS genes promoters associated with *P. aeruginosa* POA1 by psammaplin A (**1**) and its biphenylic dimer, bisaprasin (**2**).

## Results and Discussion

The marine sponge extract was partitioned between water and dichloromethane (50% v/v) using a modified Kupchan method previously described (Oluwabusola *et al*. 2020) and the CH_2_Cl_2_ fraction was further fractionated using reversed-phase solid-phase extraction (SPE). The resulting 100%SPE fraction was purified on reversed-phase HPLC to yield **1** (5.4 mg) and **2** (5.6 mg).

### 1.1 Structure Elucidation

The structure of psammaplin A (**1**)(Figure S1-S6) and bisaprasin (**2**) (Figure S7-S10) were assigned based on the interpretation of their experimental NMR and HRESIMS data and by comparison with the published data.

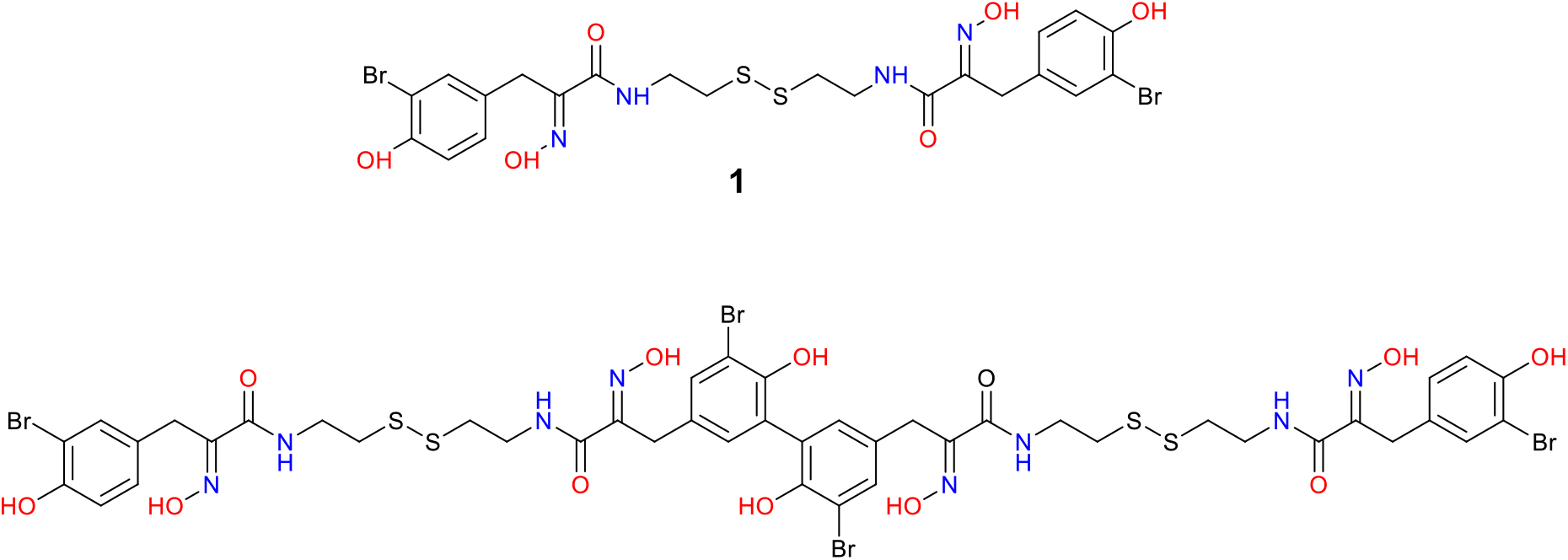

The anti-QS activities of the bromotyrosine-containing compounds, **1** and **2**, were evaluated based on biosensor strains *P. aeruginosa* POA1 quorum sensing-controlled lasB and rhlA promoters fused to an unstable gfp (ASV) reporters. The rationale behind this approach was that the growth of the bacteria in the control medium produces signal molecules at a specific threshold that induced the expression of QS-promoters genes leading to an increase fluorescence recorded every 15 min for a period of 17 hrs. For the tested compounds to show inhibition in a dose-dependent manner, it must attenuate the expression of the genes indicated by a reduced fluorescence activity.

Therefore, the dose-dependent assay against gfp-tagged *P. aeruginosa* QS-controlled *lasB* and *rhlA* promoters were determined at the concentration ranging from 100 μM to 1.563 μM. The dose-response curves of psammaplin A (**1**) and bisaprasin (**2**) when incubated with the *P. aeruginosa* PAO1 *lasB-gfp*(ASV) and *rhlA-gfp*(ASV) strains are displayed in Figure 1. The GFP expression was measured in relative fluorescence units and normalised by dividing the GFP values by the corresponding OD600 value measured at that time point.

**Figure 1.**
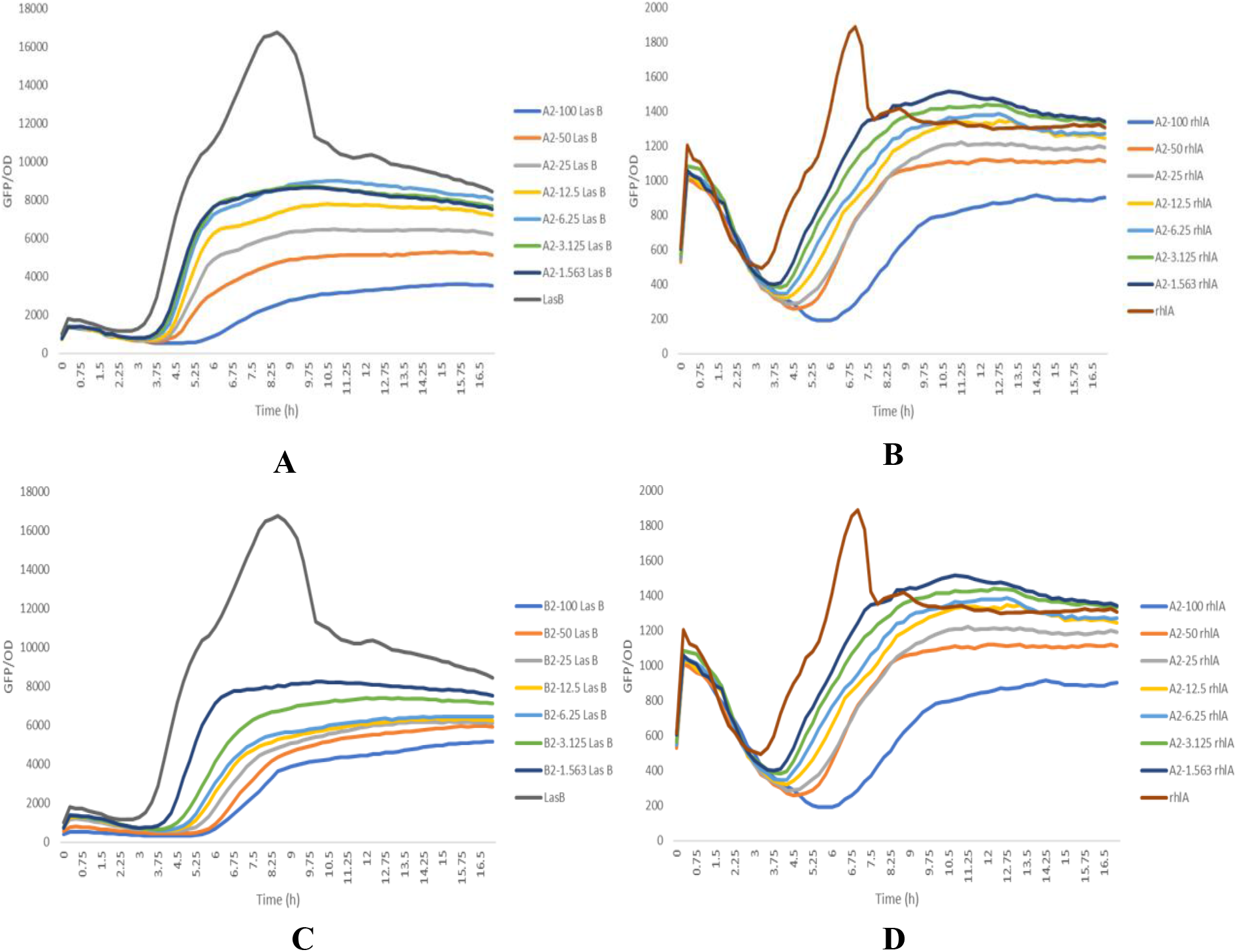
(**A**) and (**B**) are dose-response curves of psammaplin A (**1**) when incubated with *P. aeruginosa* PAO1 *LasB-gfp*(ASV) and *rhlA-gfp* (ASV) biosensor strains, respectively. (**C**) and (**D**) are dose-response curves of bisaprasin (**2**) when incubated with *P. aeruginosa* PAO1 *LasB-gfp*(ASV) and *rhlA-gfp* (ASV) biosensor strains, respectively.

The control of this experiment had the highest GFP-per-OD values, which refer to the PAO1 strain grown without the test compounds. The experiment was performed in triplicate, and the results showed significant QSIs for both compounds in a dose-dependent manner (Figure 1). The half-maximal inhibitory concentration (IC_50_ values) for the two compounds, **1** and **2**, were calculated from the dose-response curve by using Graphpad Prism 6 software package (See Figure 2). The results were obtained in a low micromolar range for **1** and **2**, showing the most significant inhibition with the IC_50_ values at 2.64 μM and 8.53 μM against the *P. aeruginosa* PAO1 *rhlA-gpf* and *lasB-gpf* biosensor strains, respectively (Table 1). By comparing the overall inhibition, **2** was more potent across the board, while **1** exhibited specific inhibition against *rhlA-gpf* biosensor strain and moderate inhibition was observed with IC_50_ value at 30.69 μM against the expression of *lasB-gpf* biosensor strain. The higher potency of **2** could be due to the dimeric nature of the molecule as compared to **1**.

**Table 1.** Quorum sensing inhibitory activity of psammaplin A and bisaprasin against *P. aeruginosa* POA1 biosensor strains.

| Compound | IC<sub>50</sub> values (μM) |  |
| --- | --- | --- |
|  | *lasB*-gpf | *rhlA*-gpf |
| 1 | 30.69 | 2.64 |
| 2 | 8.53 | 8.70 |

**Figure 2.**
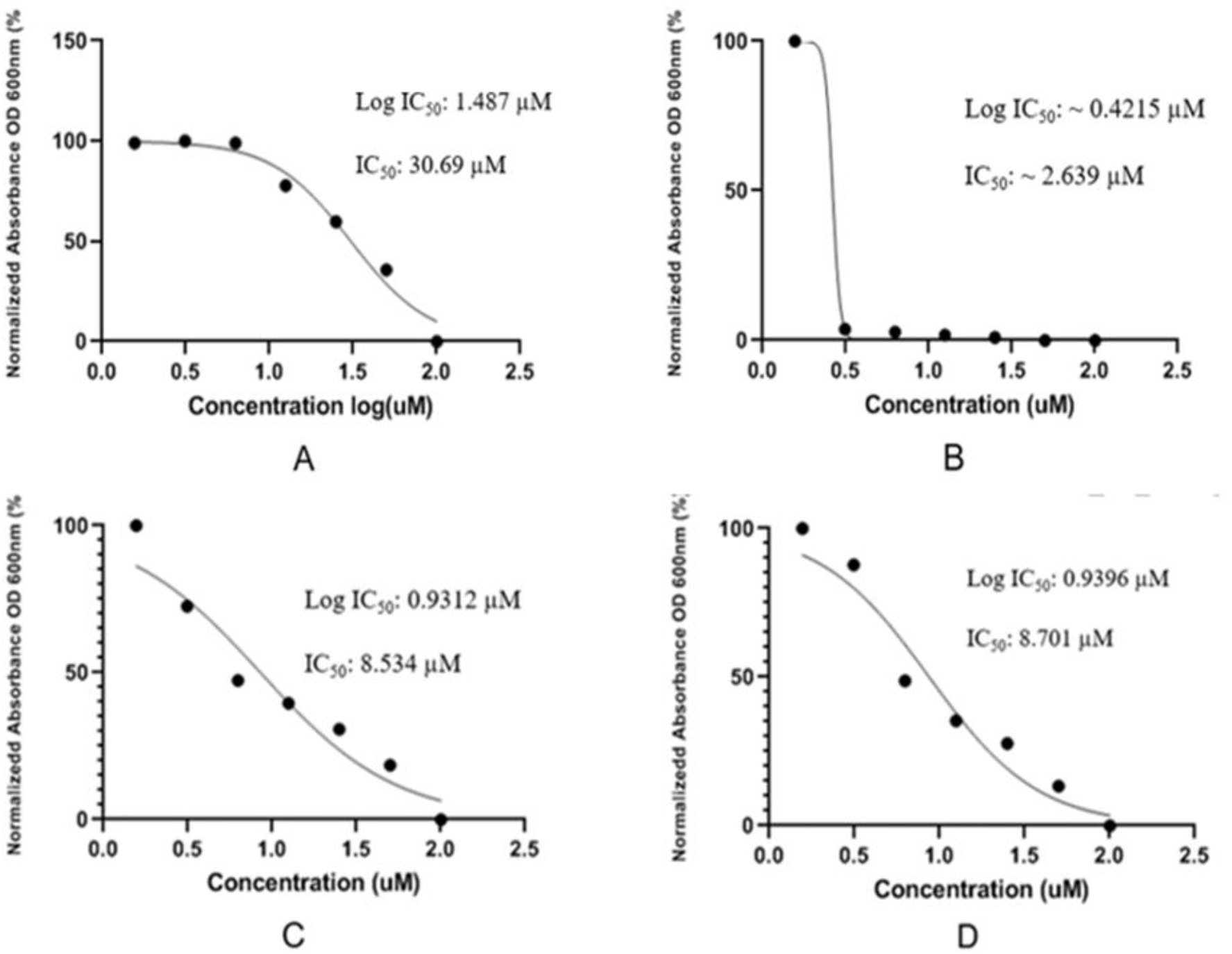
(**A**) and (**B**) are IC_50_ curves of psammaplin A (**1**) against *P. aeruginosa* PAO1 *lasB-gfp*(ASV) and *rhlA-gfp* (ASV) biosensor strains, respectively. (**C**) and (**D**) are IC_50_ curves of bisaprasin (**2**) against *P. aeruginosa* PAO1 *lasB-gfp*(ASV) and *rhlA-gfp* (ASV) biosensor strains, respectively.

The preliminary report by Davies and co-workers concluded that *las* QS system is vital in the maturation and differentiation of pathogenic bacterial biofilms as well as the expression of virulence determinant, including proteases, elastase, exotoxins and many other immune-evasion molecules (Pamp and Tolker-Nielsen 2007; Chen *et al*. 2013). Although the two brominated compounds, **1** and **2**, have not been screened against the biofilm formation of *P. aeruginosa*, it was reported that **1** and other oxime-bearing bromotyrosine derivatives, such as bastadins 3-5 and aplysamine, exhibited pronounced biofilm formation inhibition effect of *Balanus improvises* (Ortlepp *et al*. 2007). Moreso, it was established in the literature that over one hundred and thirty molecules isolated from sponges exhibited antifouling activities on diverse microorganisms or invertebrate larvae (A Database of the Marine Natural Products Literature, 2020). Based on the results obtained in this investigation together with the literature on the antibiofilm formation stated above, we have of the opinion that the mode of action of the psammaplin type compounds is likely to be interaction with the QS system of the microorganism.

## Materials and Methods

### 2.1. General Experimental Procedures

IR spectra were recorded on a PerkinElmer UATR Two, model L1600300. Both 1D and 2D NMR data were recorded on a Bruker AVANCE III HD Prodigy TCI cryoprobe at 600 and 150 MHz for ^1^H and ^13^C, respectively. HRESIMS data were obtained using a ThermoScientific LTQ XL/LTQ Orbitrap Discovery coupled to a Thermo instrument Accela HPLC system, and an Agilent 6540 HRESI-TOF-MS coupled to an Agilent 1200 HPLC system. This procedure was explained in detail in the previous publication (Oluwabusola *et al*. 2020).

### 2.2. Collection and Identification

The sponge sample was collected from the Fiji Islands in December 1997, freeze-dried and stored in 4°C. It was identified as *Aplysinella rhax* by Dr. John Hooper of the Queensland Centre for Biodiversity, Queensland Museum, Australia as described in a previous publication (Tabudravu *et al*. 2002). A voucher specimen (Voucher number: 9712SD130) is held at the Pacific Regional Herbarium at the University of the South Pacific, Suva, Fiji Islands.

### 2.3. Extraction and Isolation of Psammaplin A (**1**) and Bisaprasin (**2**)

The sponge sample was extracted with MeOH (3 × 300 mL) followed by DCM (3 × 200 mL), dried and partitioned following the modified Kupchan liquid-liquid partitioning technique described previously (Kupchan *et al*. 1978). The four fractions (*sec-*butanol fraction, methanol fraction, CH_2_Cl_2_ fraction and hexane fraction) were dried and weighed. The process of fractionation and purification that led to the isolation of compound **1** (5.4 mg) and compound **2** (5.8 mg) from the CH_2_Cl_2_ fraction was detailed in the previous publication (Oluwabusola *et al*. 2020).

### 2.4. P. aeruginosa QS. Inhibition Assays

The bromotyrosine compounds, **1** and **2**, were dissolved in 100% DMSO and mixed with ABTGC medium (AB minimal medium containing 2.5 mg/litre thiamine, supplemented with 0.2% (wt/vol) glucose and 0.2% (wt/vol) Casamino Acids), after which they were added to the first column of wells of a 96-well microtiter plate to give a final concentration of 100 μM in a final volume of 200 μL. One hundred microliters of ABTGC medium was then added to the remaining wells in the plate, and serial 2-fold dilutions of the inhibitors were made by adding 100 μL of the preceding inhibitor-containing well to the subsequent one. The final column was left without inhibitor as a control. Next, an overnight culture of the P. aeruginosa *lasB*-*gfp* and *rhlA-gfp* (ASV) strains, grown in LB medium at 37°C with shaking, were diluted to an optical density at 600 nm (OD600) of 0.2, and 100 μLl of bacterial suspension was added to each well of the microtiter plate. As such, each compound was tested at concentrations ranging from 100 μM to 1.563 μM across the plate, in a volume of 200 μL. The microtiter plate was incubated at 37°C in a Tecan Infinite 200 Pro plate reader (Tecan Group Ltd., Männedorf, Switzerland). GFP fluorescence (excitation at 485 nm, emission at 535 nm) and cell density (OD600) measurements were collected at 15-min intervals for at least 17 h.

## Acknowledgements

M.J. wishes to thank Mr Russell Gray of Marine Biodiscovery Centre, Aberdeen for running the NMR spectrometry. M. J. wishes to thank the EU. Seventh Framework Programme Project PharmaSea (grant agreement no-312184) for financial support for the collection of the marine sample. Also, This research is supported by the National Research Foundation, Prime Minister’ s Office, Singapore, under its Marine Science Research and Development Programme (award nos. MSRDP-P15 and MSRDP-P34).

## Author Contribution

J. T. collected the marine sample and M.J., R.E, and J.T. designed the study. E. T. O. undertook the extraction, isolated, purification and structure elucidation of the compounds. O. T. assisted in the extraction of the sample. L. T. T. and N. P. K. performed the biological assays and analysed the data. E. T. O. drafted the initial manuscript, which was edited and approved by all the authors.

## Supporting Information

Supplementary materials related to this article are supplied in a separate sheet.

---

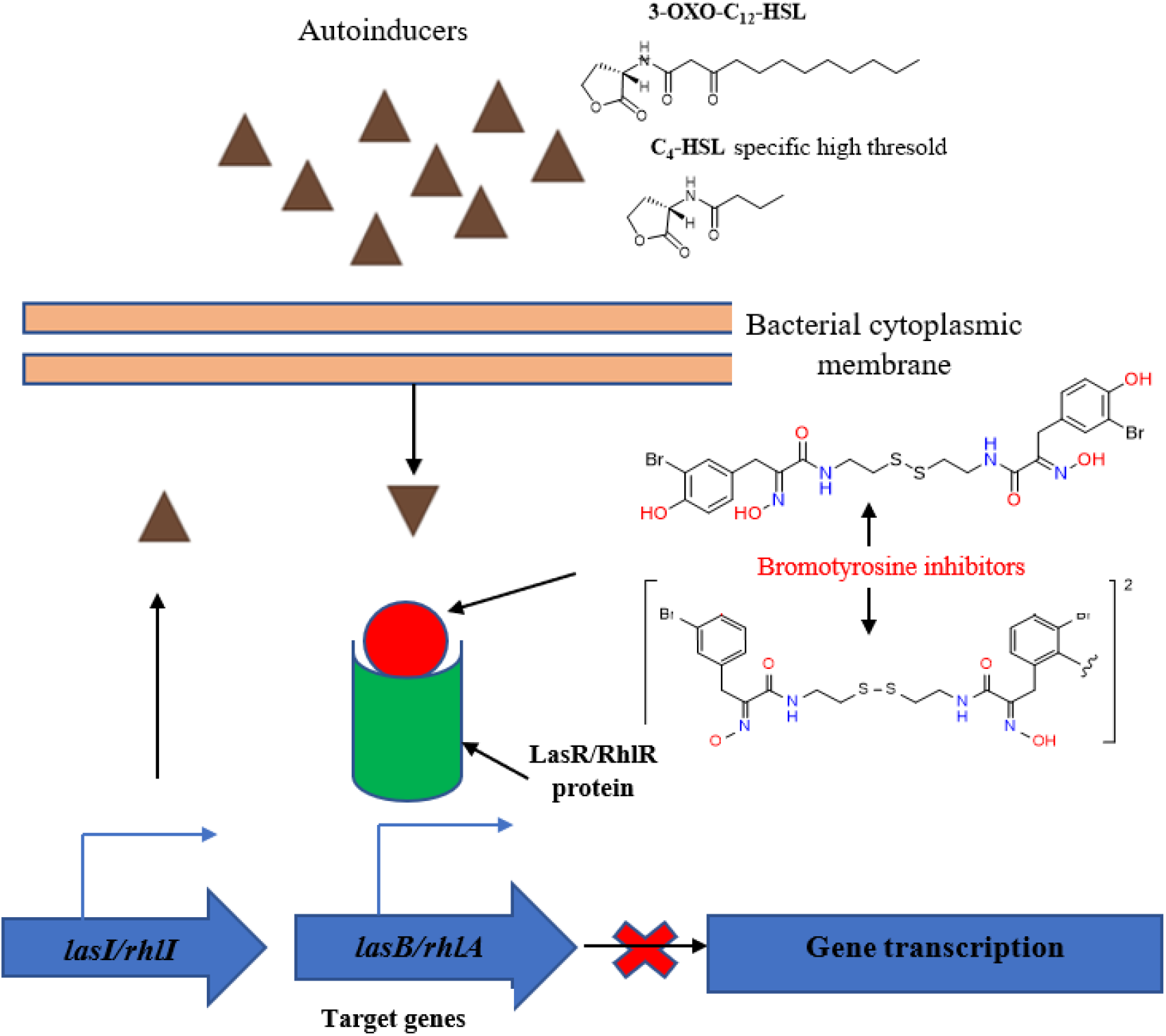

