## Supplementary material for "Inhibition of Quorum Sensing System in *Pseudomonas aeruginosa* by Psammaplin A and Bisaprasin Isolated from the Marine Sponge *Aplysinella rhax*": Supplimentary Information

**Table of content**

**Figure S1-S2** HRESIMS and ^1^H NMR spectra of **1**……………………………………………………3

**Figure S3-S4** HSQC and COSY NMR spectra of **1**……………………………………………………4

**Figure S5-S6** HMBC of **1** and Orbitrap –(+)-HRMS of **2** ……………………………………………..5

**Figure** **S7**-**S58** **^1^**H NMR and HSQC NMR spectra of **2**………………………………………………...6

**Figure** **S9**-**S10** COSY NMR and HMBC NMR spectra of **2** …………………………………………..7

**Table 1** NMR data for Psammaplin A **(1)**………………………………………………………………8

**Table 2** NMR data for Bisaprasin **(2)**…………………………………………………………………...9

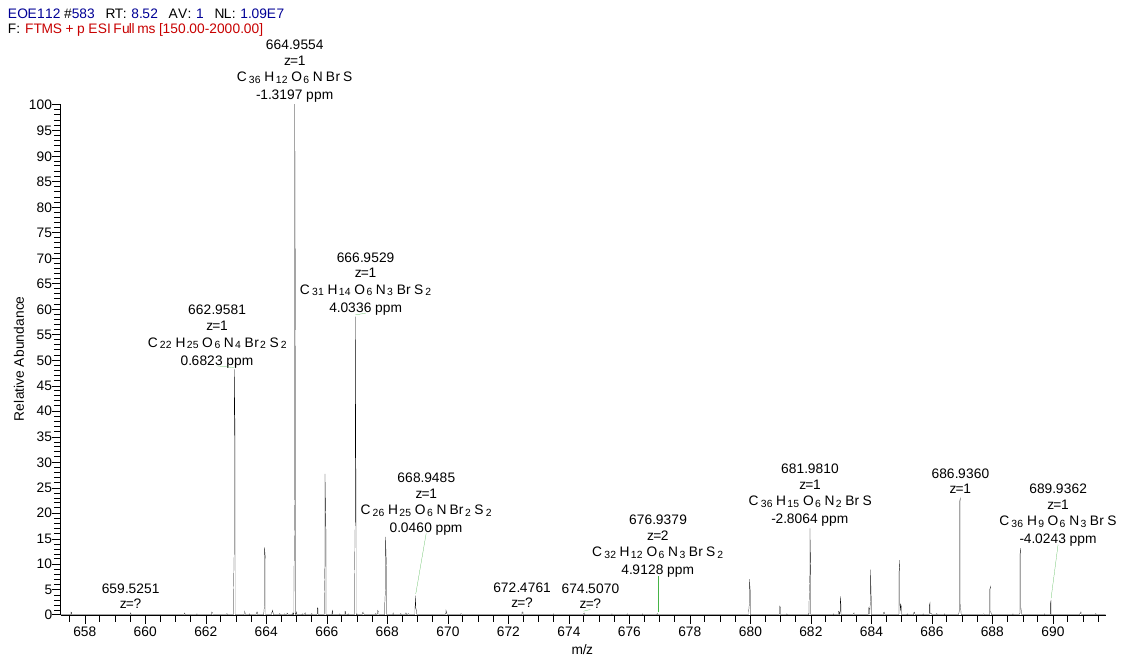

**[M+H]^+^**

**Figure S1.** Orbitrap-(+)-HRMS of **1**

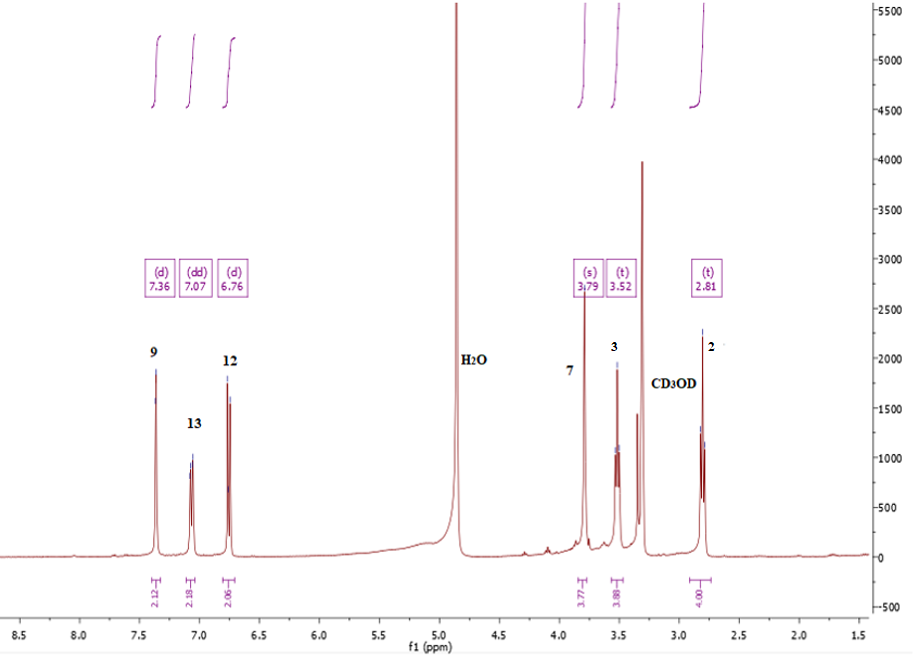

**Figure S2.** ^1^H NMR spectrum of **1** at 600MHz in CD_3_OD

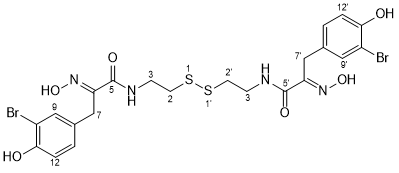

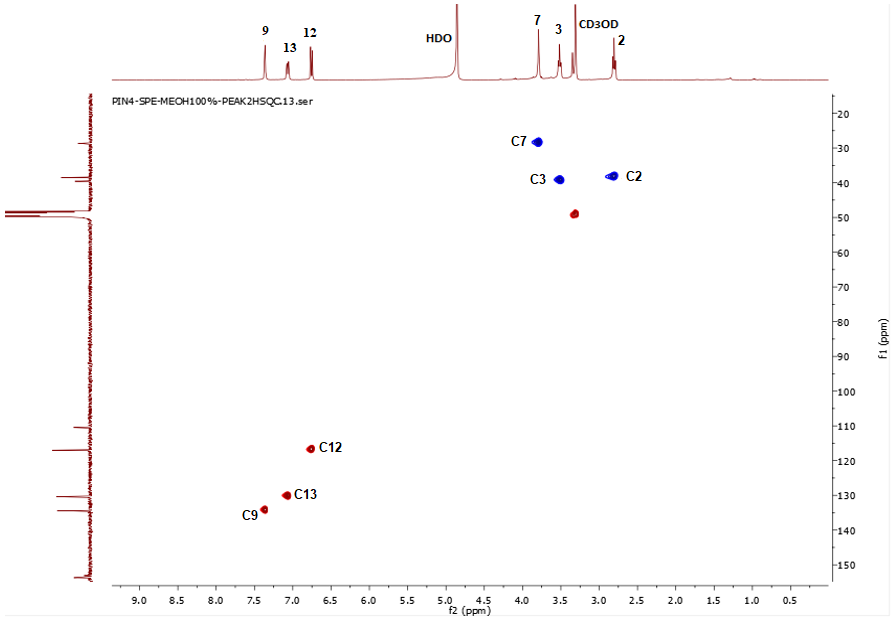

**Figure S3.** HSQC NMR spectrum of **1** at 600MHz in CD_3_OD

**Figure S4.** COSY NMR spectrum of **1** at 600MHz in CD_3_OD

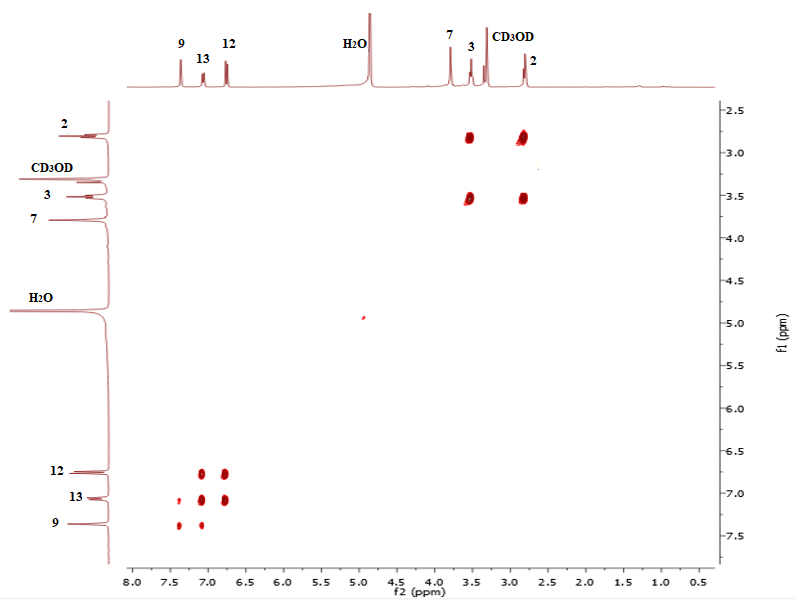

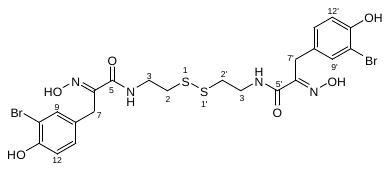

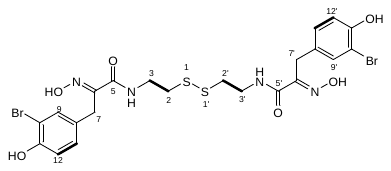

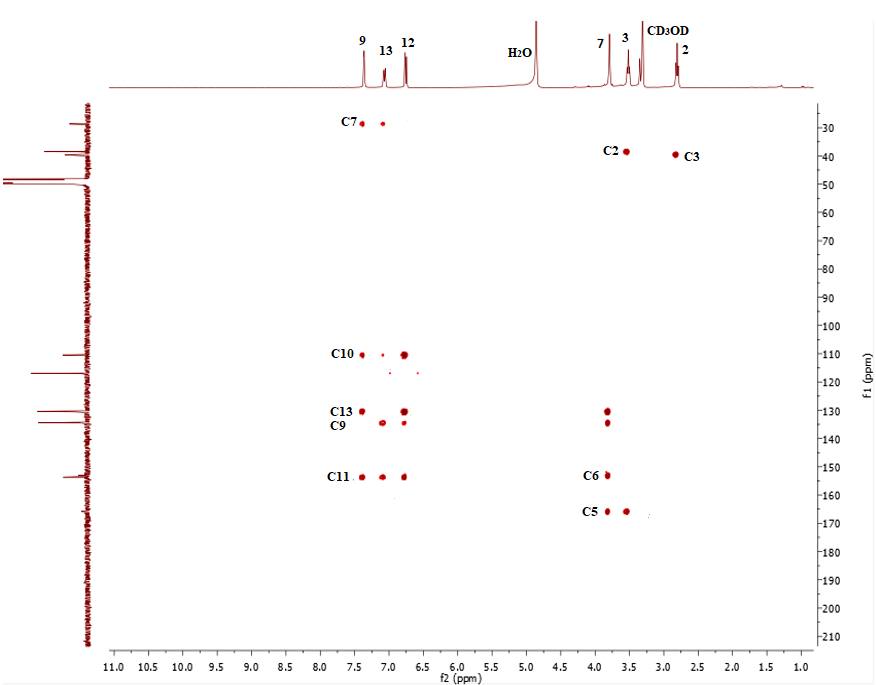

**Figure S5**. HMBC NMR spectrum of **1** at 600MHz in CD_3_OD

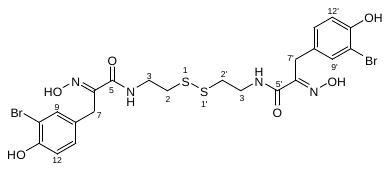

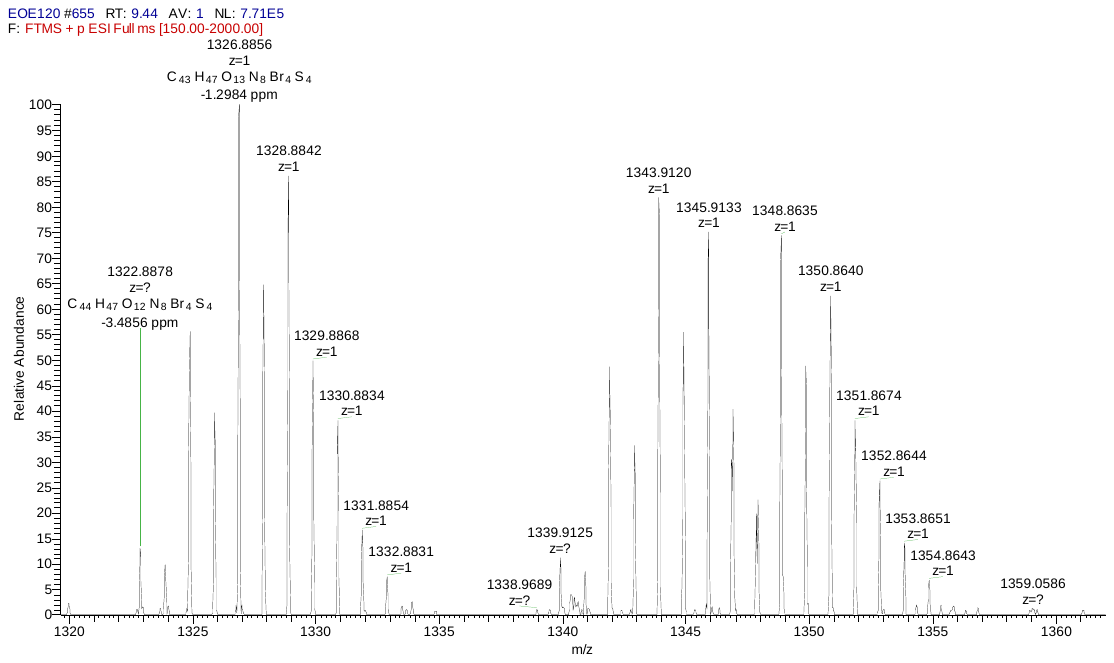

**[M+H]^+^**

**[M+NH_4_]^+^**

**Figure S6**. Orbitrap-(+)-HRMS of **2**

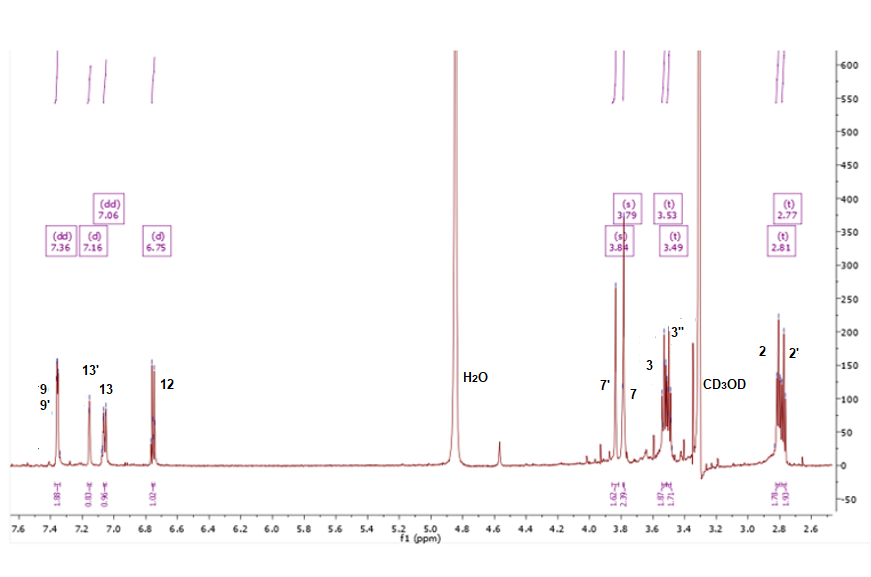

**Figure S7.** ^1^H NMR spectrum of **2** at 600MHz in CD_3_OD

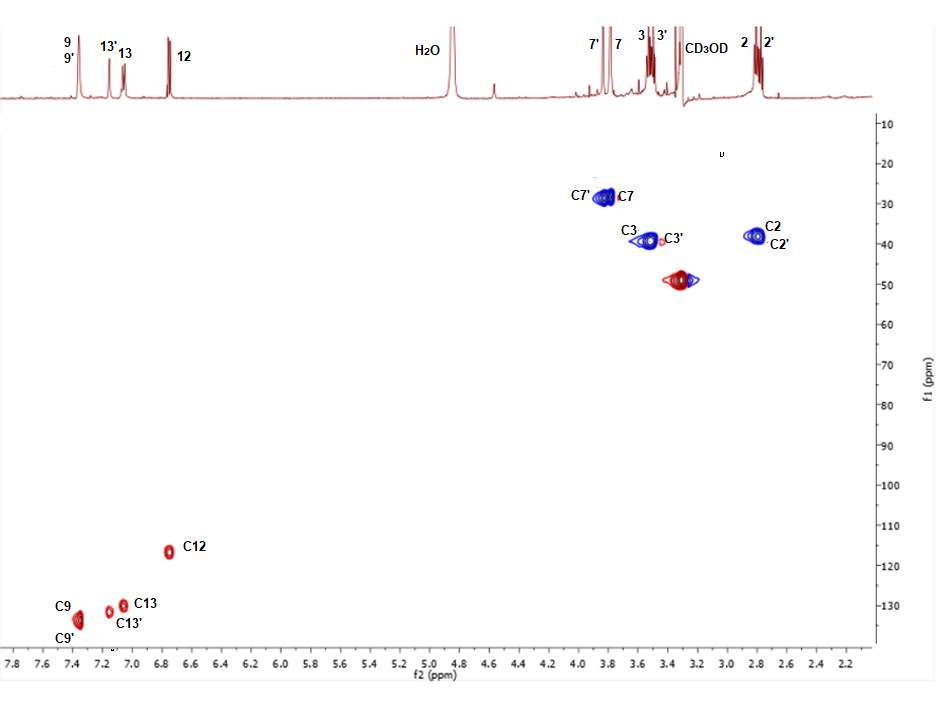

**Figure S8.** HSQC NMR spectrum of **2** at 600MHz in CD3OD

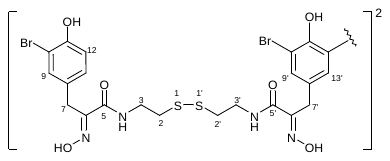

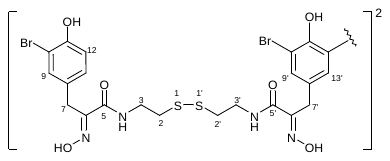

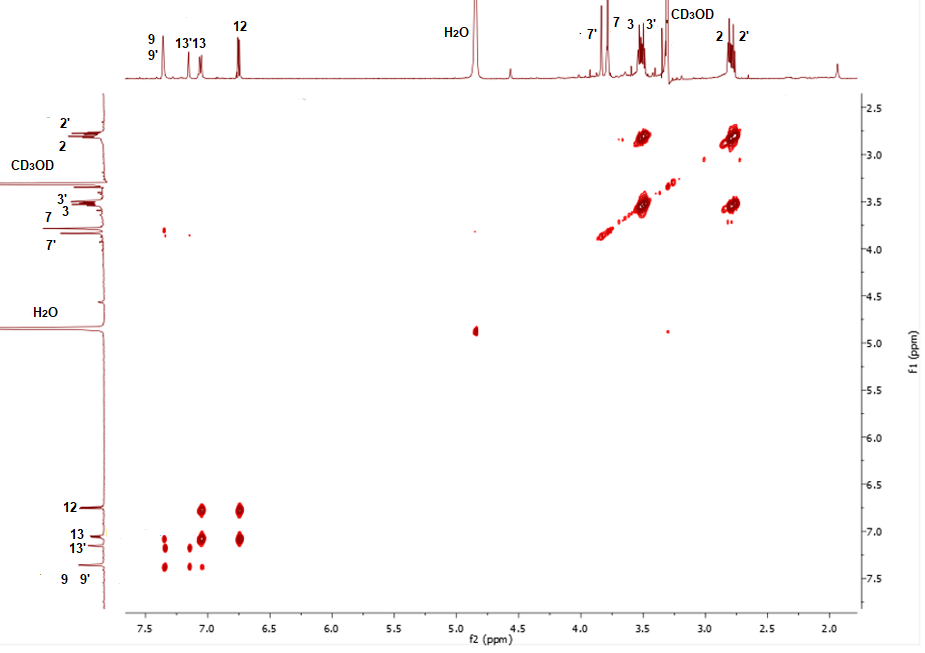

**Figure S9.** COSY NMR spectrum of **2** at 600MHz in CD_3_OD

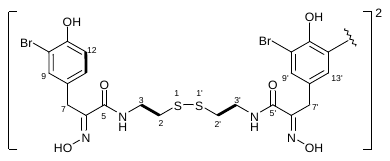

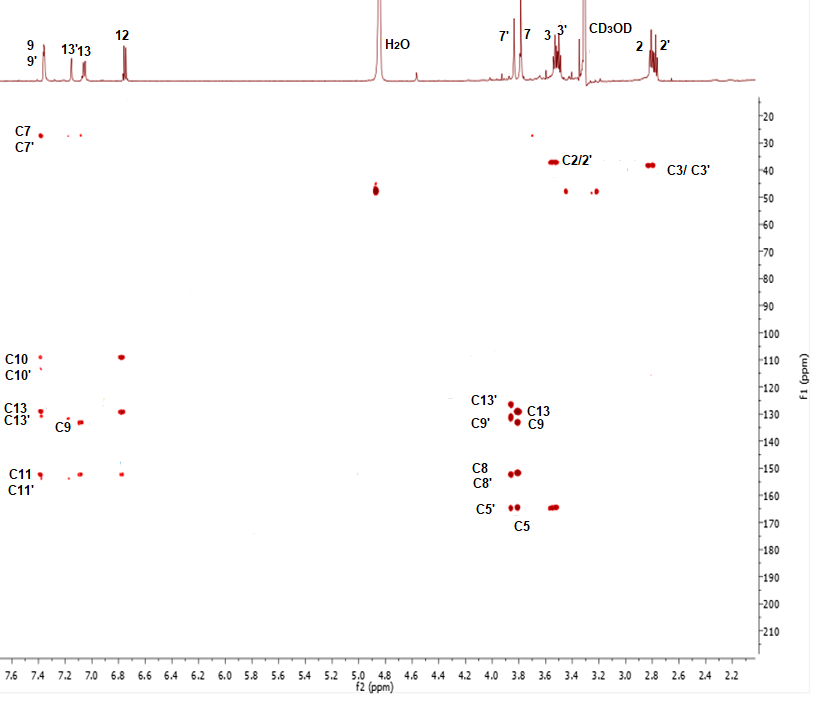

**Figure S10.** HMBC NMR spectrum of **2** at 600MHz in CD_3_OD

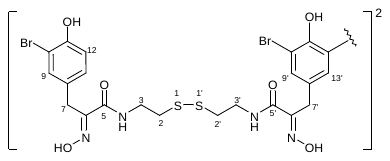

**Table 1** NMR data for Psammaplin A **(1)**

| **Position** | **δ_C,_ mult.** | **δ_H_, mult.**  **(*J* in Hz)** | **HMBC**  **(H→C)** |
| --- | --- | --- | --- |
| 2, 2’ | 38.4, CH_2_ | 2.81, t (6.0) | 3/3’ |
| 3, 3’ | 39.5, CH_2_ | 3.52, t (6.0) | 2/2’, 5/5’ |
| 5, 5’ | 165.8, C |  |  |
| 6, 6’ | 152.1, C |  |  |
| 7, 7’ | 28.6, CH_2_ | 3.79, s | 5/5’, 6/6’,9/9’ 13/13‘ |
| 8, 8’ | 130.6, C |  |  |
| 9, 9’ | 134.3, CH | 7.36, d 2.0) | 7/7’, 10/10’, 11/11’, 13/13’ |
| 10, 10’ | 110.4, C |  |  |
| 11, 11’ | 153.7, C |  |  |
| 12, 12’ | 116.8, CH | 6.76, d (8.4 ) | 9/9’, 10/10’, 11/11’, 13/13’ |
| 13, 13’ | 130.6, CH | 7.07, dd (8.4,2.0) | 7/7’, 9/9’, 10/10’, 11/11’ |
| NMR solvents used for **1** was CD_3_OD at 600 MHz. | | | |

| **Position** | **δ_C_, mult.** | **δ_H,_ mult.(*J* in Hz)** | **HMBC**  **(H → C)** |
| --- | --- | --- | --- |
| 2, 2’ | 38.4, CH_2_ | 3.52, t (6.0) | 3/3’ |
| 3, 3’ | 39.5, CH_2_ | 3.58, t (6.0) | 2/2’, 5/5’ |
| 5, 5’ | 165.8, C |  |  |
| 6, 6’ | 153.8, C |  |  |
| 7, 7’ | 28.6, CH_2_ | 3.79, s | 5/5’, 6/6’,9/9’ 13/13‘ |
| 8, 8’ | 130.6, C |  |  |
| 9, 9’ | 134.3, CH | 7.36, d (1.6) | 7/7’, 10/10’, 11/11’, 13/13’ |
| 10 , 10’ | 110.4, C |  |  |
| 11,11’ | 155.7, C |  |  |
| 12, 12’ | 116.8, CH | 6.75, d (8.0 ) | 9/9’, 10/10’, 11/11’, 13/13’ |
| 13, 13’ | 130.6, CH | 7.07, dd (8.0,1.6) | 7/7’, 9/9’, 10/10’, 11/11’ |
| 2’’,2’’’ | 38.7, CH_2_ | 2.77, t (6.0) | 3’’/3’’’ |
| 3’’,3’’’ | 39.8, CH_2_ | 3.49, t (6.0) | 2’’/2’’’, 5’’/5’’’ |
| 5’’, 5’’’ | 166.1, C |  |  |
| 6’’, 6’’’ | 153.5, C |  |  |
| 7’’, 7’’’ | 28.6, CH_2_ | 3.88 s | 5’’/5’’’, 6’’/6’’’, 9’’/9’’’, 13’’/13’’’ |
| 8’’, 8’’’ | 128.3, C |  |  |
| 9’’, 9’’’ | 133.4, CH | 7.36, d (1.6) | 7’’/7’’’, 10’’/10’’’, 11’’/11’’’, 13’’/13’’’ |
| 10’’ , 10’’’ | 111.4, C |  |  |
| 11’’,11’’’ | 153.1, C |  |  |
| 12’’, 12’’’ | 112.7, C |  |  |
| 13’’, 13’’’ | 130.5, CH | 7.16, d (1.6) | 7’’/7’’’, 9’’/9’’’, 10’’/10’’’, 11’’/11’’’ |
| NMR solvents used for **2** was CD_3_OD at 600 MHz. | | | |

**Table 2** NMR data for Bisaprasin **(2)**
